# Functional effects of heating and cooling gene networks

**DOI:** 10.1101/181537

**Authors:** Daniel A. Charlebois, Kevin Hauser, Sylvia Marshall, Gábor Balázsi

## Abstract

Everyday existence and survival of most organisms requires coping with temperature changes, which involves gene regulatory networks both as subjects and agents of cellular protection. Yet, how temperature affects gene network function remains unclear, partly because natural gene networks are complex and incompletely characterized. Here, we study how heating and cooling affect the function of single genes and well-characterized synthetic gene circuits in *Saccharomyces cerevisiae*. We found nontrivial, nonmonotone temperature-dependent gene expression trends at non-growth-optimal temperatures. In addition, heating caused unusual bimodality in the negative-feedback gene circuit expression and shifts upward the bimodal regime for the positive feedback gene circuit. Mathematical models incorporating temperature-dependent growth rates and Arrhenius scaling of reaction rates captured the effects of cooling, but not those of heating. Molecular dynamics simulations revealed that heating alters the conformational dynamics and allows DNA-binding of the TetR transcriptional repressor, fully explaining the experimental results for the negative-feedback gene circuit. Overall, we uncover how temperature shifts may corrupt gene networks, which may aid future designs of temperature-robust synthetic gene circuits.

## 1 Introduction

Temperature plays a critical role across all of biology. Endotherm organisms, commonly called “warm-blooded”, maintain their body temperature and perish if this regulation fails. Despite common portrayal of most species as “cold-blooded”, even ectotherms and unicellular microbes prefer a specific temperature range where they function optimally. If the ambient temperature departs from this optimum, cells and organisms attempt to minimize the potential damage. For instance, the heat-shock response elicits the transient expression of cytoprotective proteins, mitigating a multitude of harmful effects ranging from changes in cell morphology to alteration in protein structure and function (Richter et al, 2010; Verghesea et al, 2012). Lack or failure of heat-shock response leads to prolonged cell cycle arrest (Yost & Lindquist, 1986; Zeuthen, 1971), or even cell death.

Gene regulatory networks govern cellular protein levels and play a double role during temperature changes. First, they require protection due to their crucial role in cellular housekeeping and homeostasis (Watson et al, 2013), development (Levine & Davidson, 2005), and survival (Charlebois et al, 2014). Second, they are also instrumental in generating the protective stress response. This twofold involvement as both the subject and agent of cellular protection makes it difficult to understand what happens with gene networks after temperature shifts occur. Additional complexity stems from nontrivial effects of temperature on transcription (Oliveira et al, 2016), translation (Neupert et al, 2008) and their regulation (Madrid et al, 2002; Perez-Martin & Espinosa, 1994). Previous transcriptome expression measurements in heat and cold shock revealed broad responses (Gasch et al, 2000), but it is hard to discern the degree to which these changes were individual gene-(Arnaud et al, 2015) or network-level effects, or cellular attempts to restore homeostasis. Adding to these difficulties are the complexity of biochemical interaction networks and the incomplete knowledge of their connectivity (Charlebois et al, 2007; Yu et al, 2008).

Synthetic gene circuits are relatively simple human-designed gene networks that perform specific predefined functions, and have promising roles as switches (Gardner et al, 2000), oscillators (Elowitz & Leibler, 2000; Stricker et al, 2008), and logic gates (Anderson et al, 2007) in future medical (Ro et al, 2006; Saxena et al, 2016; Slomovic et al, 2015), bioenergetic (Dunlop et al, 2011), or environmental (Windbichler et al, 2011) applications. The connectivity and components of synthetic gene circuits are typically well-characterized and completely known by design, and their regulatory connections with the host network are assumed minimal. Therefore, these relatively simple gene network modules should be less affected by cellular attempts to restore homeostasis, making them viable candidates for discerning and separately studying the network-level effects of temperature changes. Besides a potential to advance fundamental understanding of temperature effects on biological systems, future synthetic biology applications will require accuracy and robustness in carrying out a predefined function, independent of environmental changes (Cardinale & Arkin, 2012; Zechnera et al, 2016). In some cases temperature provides a way to control synthetic gene circuits (Isaacs et al, 2003) and temperature compensation may be possible (Hussain et al, 2014). However, since most synthetic gene circuits are developed and characterized under controlled laboratory conditions, it is unclear how temperature shifts affect their function. Overall, by quantitatively and predictively studying how synthetic gene circuits respond to heating and cooling, we might also learn how temperature affects natural gene networks.

To address these questions, we chose to study two single genes and two synthetic gene circuits with autoregulation. Positive and negative autoregulation are highly abundant motifs in natural gene networks (Ferrell, 2013; Rosenfeld et al, 2002; Thieffry et al, 1998), and have been commonly used in synthetic biology (Atkinson et al, 2003; Basu et al, 2004; Becskei & Serrano, 2000). The first synthetic biological construct we employ is the ‘linearizer’ or Negative Feedback (NF) gene circuit integrated into the budding yeast genome (Nevozhay et al, 2009). It consists of the tetracycline repressor (TetR) inhibiting transcription of its own gene and a reporter gene from identical P_GAL1-D12_ promoters in an inducer-dependent manner (Blake et al, 2006; Murphy et al, 2007). This gene circuit reduces the heterogeneity of gene expression at a wide range of inducer concentrations, and linearizes the dose response prior to saturation compared to similar gene circuits without autoregulation (Nevozhay et al, 2009). The second construct we study is the positive feedback (PF) gene circuit (Nevozhay et al, 2012), originally used to investigate how the costs and benefits of gene expression shape population dynamics and thereby genetic evolution (González et al, 2015). The PF gene circuit consists of an rtTA transactivator that activates its own gene and a reporter gene when bound by a tetracycline analog inducer (Nevozhay et al, 2012). Since the PF and NF gene circuit components are not native to yeast and lack substantial connections with the native yeast regulatory network, their behavior at 30°C was predictable by quantitative modeling (Nevozhay et al, 2009; Nevozhay et al, 2012). This raises the possibility that by studying the NF and PF gene circuits in relative isolation from temperature compensation and other natural network complexities, we may unravel how heating and cooling affect gene network function.

To unravel how temperature affects gene circuit function, we measured both gene expression and cell division rates, which mutually influence each other (Klumpp et al, 2009; Nevozhay et al, 2012; Scott et al, 2010; Tan et al, 2009). Every cell type we investigated grew fastest at 30°C, a possible consequence of culturing in standard laboratory conditions. We found nontrivial temperature-dependent gene expression trends for single reporter genes. These dependencies became more complex for induced gene circuits.

Computational models with temperature-dependent growth rates and Arrhenius reaction kinetics captured the functional effects of cooling, but not those of heating. On the other hand, molecular dynamics (MD) simulations revealed a novel ability of inducer-bound, normally inactive TetR to repress its target promoter. Once we augmented the quantitative models with this altered protein behavior, we could fully explain the changes in gene circuit function at all temperatures. Overall, these results suggest ways to understand and predict how heating and cooling affect gene network function, and warn about the unexpected effects arising from altered repressor conformational dynamics at high temperatures.

## 2 Results

### 2.1. Temperature-dependent growth and expression in cells with a single reporter gene

Since regulatory networks consist of genes, we began by investigating how temperature affects single reporter genes. Therefore, we focused on PF0 and NF0 cells that carried only a reporter gene, but lacked the corresponding regulators (see the Methods and Fig. 1A,B). Specifically, the PF0 strain lacked the rtTA activator, but contained the unregulated yEGFP::zeoR reporter gene driven by the *pCYC1*-derived P_TETREG_ promoter (Fig. 1A). Likewise, the NF0 strain lacked the tetR repressor, but contained the same reporter gene driven by the *pGAL1*-based P_GAL1-D12_ promoter (Fig. 1B), which maintains natural function despite two *TetO2* operator sites inserted behind the TATA box (Blake et al, 2006; Blake et al, 2003).

**Figure 1.**
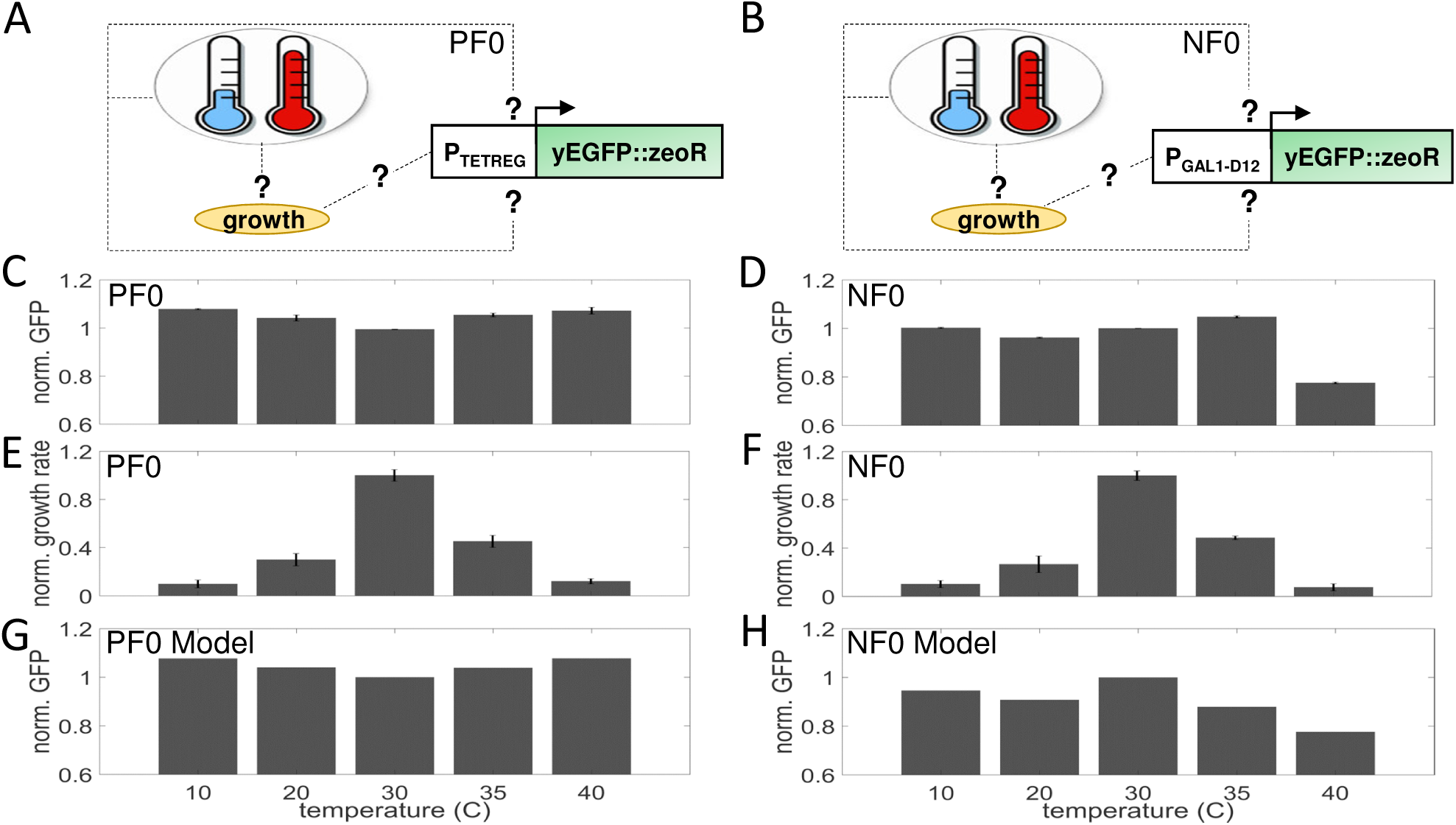
The effect of temperature on single reporter strain fitness and gene expression. **(A)** Schematics of PF0 consisting of the yEGFP::zeoR reporter controlled by the P_TETREG_ promoter. **(B)** Schematics of NF0 consisting of the yEGFP::zeoR reporter controlled by the P_GAL1-D12_ promoter. **(C)** PF0 population average (mean) reporter expression as a function of temperature. **(D)** NF0 population average (mean) reporter expression as a function of temperature. **(E)** Fitness (growth rate) of PF0 cells as a function of temperature. **(F)** Fitness (growth rate) of NF0 cells as a function of temperature. **(G)** Gene expression results from Arrhenius models for PF0 cells (see Table S2 for parameters). **(H)** Gene expression results from Arrhenius models for NF0 cells (see Table S2 for parameters). Fitness and GFP values were normalized by the corresponding values from replicates in the 30°C control condition (C-H). Error bars are SEM (*N* = 3).

Varying the temperature from 10°C to 40°C, we observed convex temperature-dependence of gene expression for PF0 cells (Figs. 1C and S1), with a minimum around 30°C. On the other hand, the gene expression trend for NF0 cells was more complex, with a peak at 35°C (Figs. 1D and S1). Considering that cellular growth rates and gene expression mutually affect each other (Klumpp et al, 2009; Nevozhay et al, 2012; Tan et al, 2009), we asked if the temperature-dependence of gene expression could arise from growth rate differences. Therefore, we measured the growth rates of NF0 and PF0 cells at the same temperatures as their expression. The growth rates of PF0 and NF0 cells were indeed temperature-dependent, with optima at 30°C (Figs. 1E,F and S1A,B). In contrast to earlier results in bacteria (Ratkowsky et al, 1982), the temperature dependence (*R*^2^ > 0.98) of both PF0 and NF0 growth rates followed the Arrhenius law below the growth optimum at 30°C (Fig. S2A,B). Above the optimum, growth rates declined rapidly, approximately linearly with temperature.

To examine if these experimental temperature-dependent trends of gene expression were purely due to growth rate changes, we developed computational “growth rate models” for single genes, setting all protein dilution rates equal to the experimental temperature-dependent growth rates (see section SI 1). These growth rate models could only reproduce the experimentally observed trends for the PF0 strain qualitatively (Fig. S3A,B). Therefore, next we also assumed that protein synthesis rates had Arrhenius-type temperature dependence. These augmented “Arrhenius models” quantitatively matched the gene expression trend observed in PF0 cells (Fig. 1G), and also NF0 cells, except at 35°C (Fig. 1H). This discrepancy at 35°C could arise because the P_GAL1-D12_ promoter is native P_GAL1_ promoter-based, with sugar-dependent regulation that may be temperature-sensitive. On the other hand, the more generically (aerobically) active P_CYC1_ promoter (Burke et al, 1997), from which the P_TETREG_ promoter was derived, matches better the constitutively expressed gene assumption in the models. We concluded that including temperature-dependent growth and reaction rates into Arrhenius models was sufficient to capture temperature-dependent expression changes for a single reporter gene, especially from a bona-fide constitutive promoter.

### 2.2 Temperature-dependent growth and expression in uninduced PF and NF cells

The PF and NF gene circuits are simple examples of hierarchical networks, in which environmental effects can percolate down through regulatory layers, increasing the difficulty of their interpretation. For simplicity, we started by studying the temperature-dependence of their expression without inducer. The single-gene measurements guided us to first study the exponential growth rates of budding yeast cells with uninduced PF (Fig. 2A) and NF (Fig. 2B) gene circuits ( “PF cells” and “NF cells” hereafter, respectively) at constant temperatures ranging from 5°C to 40°C (see the Methods). Both uninduced PF and NF cells grew optimally at 30°C, and their growth rates decreased similarly (Figs. 2C,D and S4A,B) to PF0 and NF0 strains at other temperatures; a possible effect from decades-long growth in standard laboratory conditions.

**Figure 2.**
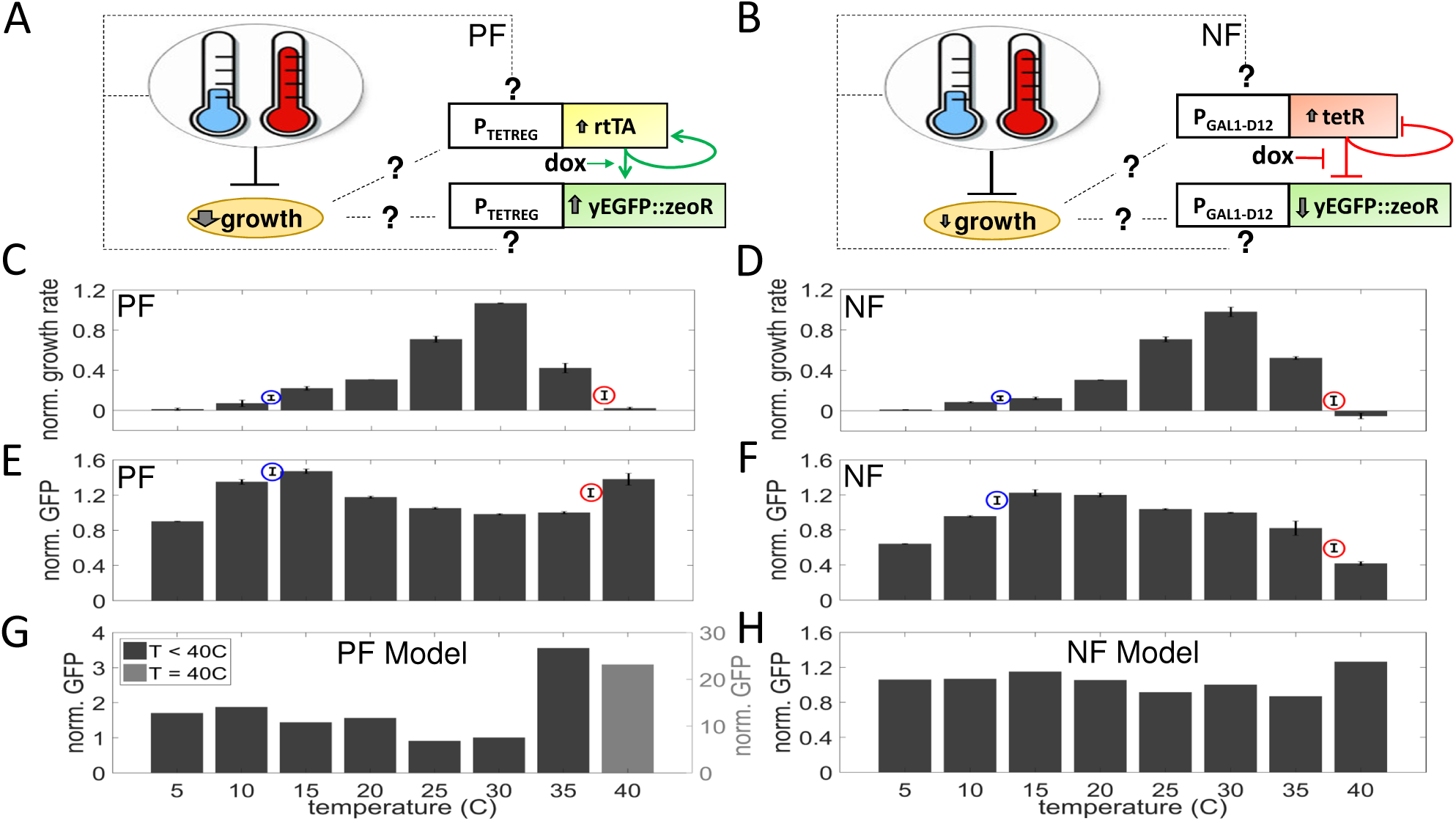
The effect of temperature on uninduced PF and NF cell fitness and gene expression. **(A)** Schematic of positive feedback (PF) synthetic gene circuit, consisting of the transactivator rtTA that activates itself and the yEGFP::zeoR reporter when bound by the inducer doxycycline (dox). Pointed arrows indicate activation. **(B)** Schematic of the negative feedback (NF) synthetic gene circuit, consisting of the yEGFP::zeoR reporter and the tetR repressor that also regulates its own expression. Repression is relieved by dox. blunt arrows indicate repression. **(C)** Fitness (growth rate) of uninduced PF cells as a function of temperature. **(D)** Fitness (growth rate) of uninduced NF cells as a function of temperature. **(E)** Average (mean) reporter expression as a function of temperature for uninduced PF populations. **(F)** Average (mean) reporter expression as a function of temperature for uninduced NF populations. **(G)** Temperature dependent gene expression results from Arrhenius models for PF cells. (See Table S2 for parameters). GFP at 40°C was plotted on a separate scale (grey bar). **(H)** Temperature dependent gene expression results from Arrhenius models for NF cells. (See Table S2 for parameters). Blue and red circles indicate mean growth rates (C and D) and GFP levels (E and F) at 12°C and 38°C, respectively. Fitness and GFP values were normalized by the corresponding values from replicates in the 30°C control condition (C-H). Error bars are SEM (*N* = 3).

Next, we studied the temperature-dependence of basal expression for intact but uninduced NF and PF gene circuits at the same temperatures as the growth measurements (see the Methods). Since all gene expression distributions were unimodal (Fig. S5), we plotted their temperature-dependent averages as expression surrogates in Fig. 2E,F. As opposed to the growth rates, we observed nontrivial, nonmonotone temperature-dependent trends of gene expression for both PF and NF cells. Specifically, the average gene expression in PF cells first increased as temperatures deviated from 30°C, but then it decreased for temperatures below 15°C (Figs. 2E and S4C). On the other hand, the gene expression of NF cells seemed to have a global peak at ∼15°C, and then decreased with varying slopes as a function of temperature (Figs. 2F and S4D).

Interestingly, temperature-effects on gene expression in uninduced PF cells were more pronounced (Fig. 2E), but otherwise very similar to those in PF0 cells (Fig. 1C), with a minimum at the standard temperature of 30°C. In contrast, the temperature-dependences of the NF0 strain (Fig. 1D) and the uninduced NF strain (Fig 2F) differed substantially, especially at low temperatures where the NF0 strain did not peak. Yet, the sharp drop in gene expression above 35°C seemed to be preserved even between these strains. We reasoned that the similar temperature-dependencies of PF0 and uninduced PF cells may relate to lack of regulation in uninduced PF. Indeed, only inducer-bound rtTA can activate its target genes (Fig. 2A), so reporter gene expression should be independent of rtTA in uninduced PF cells, similar to PF0 (Figs. 1C,E and 2C,E). In contrast, TetR repression is fully present in uninduced NF cells, which may explain their difference from NF0 cells. We reasoned that regulation should cease instead in fully induced NF cells (Fig. 2B), resembling NF0. Indeed, expression dropped similarly in both NF0 (Fig. 1D) and fully induced NF cells (Fig. S6A) at low and high temperatures, as expected.

To understand the temperature effects on uninduced NF and PF cells, we used the same two types of computational models (growth rate and Arrhenius models) as above. First, incorporating the growth rates of these strains (Fig. S4A,B) into previously established ordinary differential equation (ODE) models (Nevozhay et al, 2009; Nevozhay et al, 2012) of the NF and PF gene circuits (Eqns. 2 and 3, respectively) reproduced qualitatively the concave temperature-dependence of uninduced PF cells between 15°C and 40°C, but failed to match the gene expressions trend for uninduced NF cells (Fig. S3C,D). Next, we additionally incorporated temperature-dependent reaction rates to obtain Arrhenius models (see section SI 2). These Arrhenius models matched the experimental data reasonably well (Figs. 2G,H), except for the large increase in gene expression at 40°C, which arose computationally from rapidly falling growth (dilution) rates (Fig. 2A,B and S2C,D) and rising synthesis rates of proteins assumed to still fold properly.

In summary, incorporating temperature-dependent growth and reaction rates into Arrhenius models of constitutive gene expression could reproduce the effect of cooling, but could only partially account for the effect of heating on gene expression, especially for NF cells. Considering their better performance, subsequently we only used Arrhenius models. Also, based on the measurements of single genes and uninduced gene circuits, we selected 12°C as the “cold” and 38°C as the “warm” temperatures to study further, compared to the “standard” 30°C condition. These temperatures slowed, but did not halt cell growth (Fig. S7-S9), so that we could still dilute and passage the cell cultures into fresh medium. Earlier evidence that most yeast proteins remain folded below 40°C (Dill et al, 2011) and above 5°C (Sawle & Ghosh, 2011) further justified these growth temperature choices.

### 2.3 Temperature-dependent growth rates of induced PF and NF cells

Armed with an understanding how temperature affects cell growth and expression of single genes, uninduced or fully induced networks, next we asked how temperature affects network behaviors if we also adjust the strength of regulatory connections by altering the inducer level. However, these effects are more difficult to unravel because gene circuit induction can curb growth rates even at standard temperature (Nevozhay et al, 2012). The inducer must impede growth only indirectly, through the gene circuits since doxycycline did not alter the fitness of “parental” YPH500 yeast cells devoid of synthetic gene circuit components (Fig. S8).Therefore, we first asked how temperature interacts with induction to alter PF and NF cellular growth rates.

To address this question, we measured fitness (defined as the exponential growth rate) of PF and NF cells at various inducer (doxycycline) concentrations at warm, cold, and standard temperatures (Fig. 3A,B). We observed that warm and cold PF and NF cells grew much slower than at 30°C throughout all induction levels (Figs. 3A,B and S7A,B). Therefore, nonstandard temperatures strongly reduce the growth of PF and NF cells at all inducer concentrations. Nonetheless, induction further lowers the fitness of cells with gene circuits (Fig. 3A,B). For PF cells (Nevozhay et al, 2012) this fitness decline stems from activator (rtTA) “squelching” (Ptashne & Gann, 1990) by general transcription factor sequestration from vital cellular processes (Baron et al, 1997). To separate the effects of temperature and doxycycline-induced gene expression on fitness, we normalized the growth rates in each temperature by their corresponding uninduced values (doxycycline = 0 μg/ml; Figs. S9 and S7C,D). Such normalization eliminates temperature-related effects, so any remaining differences relative to standard temperature should indicate an interaction between induction and temperature. Interestingly, the normalization revealed that low induction still decelerated the growth of warm and cold PF cells, but less than at 30°C (Fig. S9A). This suggests that both heating and cooling might lessen inducer toxicity at low induction. On the other hand, at high induction heating seems to worsen induction toxicity (Fig. S9A), possibly as spurious rtTA interaction affinities towards general transcription factors increase in Arrhenius-manner.

**Figure 3.**
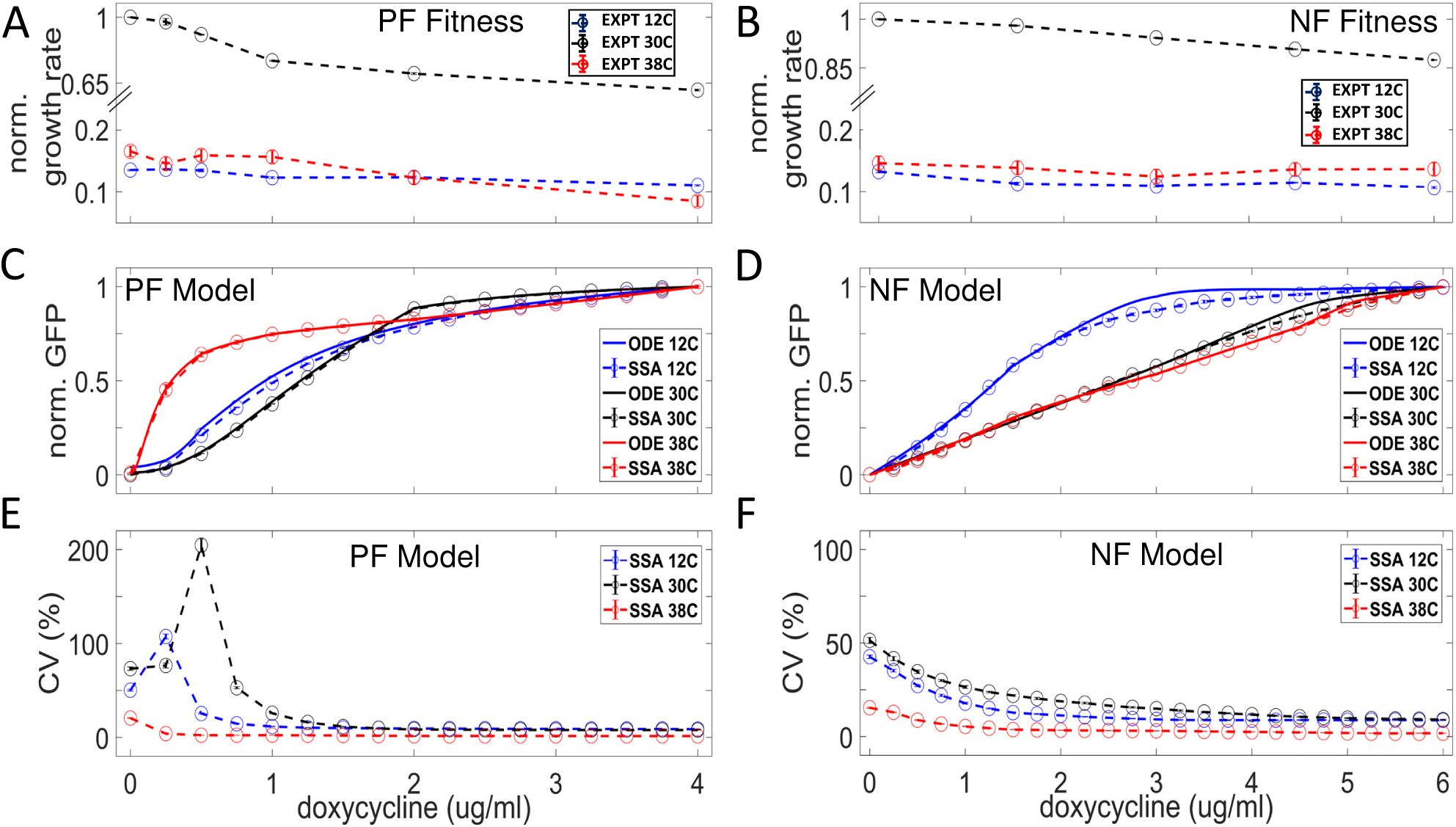
Fitness and temperature-mediated dose-response changes predicted from Arrhenius models. **(A)** Experimentally determined fitness (growth rate) for PF cells as a function of varying inducer (doxycycline) concentrations at 12°C, 30°C, and 38°C. **(A) (B)** Experimentally determined fitness (growth rate) for NF cells as a function of varying inducer (doxycycline) concentrations at 12°C, 30°C, and 38°C. **(C)** Models of mean gene expression for PF cells. **(D)** Models of mean gene expression for NF cells. **(E)** Modeling results for coefficient of variation (CV) of gene expression for PF cells. **(F)** Modeling results for coefficient of variation (CV) of gene expression for NF cells. Parameters for (C)-(F) for non-control (12°C and 38°C) conditions were adjusted according to the Arrhenius equation, where all activation energies were constrained to be between 55 kJ/mol and 87 kJ/mol (see Methods and SI for details). Error bars are SEM (*N* = 3). See Table S2 for parameters.

Whereas the fitness of NF cells was much less inducer-dependent than that of PF cells in all temperatures, some opposite trends to PF were noticeable. The same normalization revealed that heating and cooling increased the slight toxicity of moderate NF induction (Fig. S9B). On the other hand, high NF induction toxicity seemed to vanish in warm conditions (Fig. S9B). Unlike for PF, the inducer-dependent fitness decline in NF cells likely originates from transcription and translation costs alone (Kudla et al, 2009; Scott et al, 2010), which seem to depend differently on temperature than squelching.

### 2.4 Temperature effects on the dose responses of PF and NF cells

Next, we asked if quantitative models of NF and PF cells could predict how temperature affects the dose-response curves of gene expression versus inducer concentrations based on the knowledge we gained so far. To make such predictions, we developed Arrhenius versions of previous models of these gene circuits (Nevozhay et al, 2009; Nevozhay et al, 2012) and fit them to gene expression data in the standard temperature condition (see section 4.1 in Materials and Methods, and section 2 in SI Appendix for details). Unlike for uninduced gene circuits, some histograms were broad or bimodal at nonzero induction levels. Therefore, we also studied the temperature-dependence of the coefficient of variation (CV, defined as the ratio of the standard deviation to the mean) to characterize the widths of the distributions, in addition to the mean gene expression at various inducer levels.

The computational models predicted that cooling should increase the inducer-sensitivity of both PF and NF gene expression dose-responses relative to the standard temperature (Figs. 3C,D and S10A,C). Additionally, heated PF cells should be even more inducer-sensitive than cooled PF cells (Fig. 3C). For the NF gene circuit, we predicted that heating should not affect the gene expression dose response (Fig. 3D). Moreover, the models predicted that cooling should shift the PF CV peak leftward, toward lower inducer concentrations (Fig. 3E), while the CV peak of heated PF cells should vanish. We also predicted that cooling should leave the peakless NF CV dose-response unaffected, while heating should decrease it (Fig. 3F).

To test these computational predictions, we obtained reporter expression distributions at various inducer levels for the PF gene circuit. Heating and cooling increased the inducer sensitivity of the mean pf dose response (Fig. 4A) as predicted computationally, although more strongly (Fig. 3C). All PF CV dose-responses had a peak. The position and height of this peak increased with the temperature (Fig. 4B), corresponding to the emergence of bimodal PF histograms (Fig. 4C-E). This contradicted the peakless CV curve predicted computationally for heated PF cells (Fig. 3E and S10B). Overall, temperature seemed to reduce the inducer-sensitivity and enhance the expression noise of the PF gene circuit, suggesting a temperature-dependent decline in rtTA activity.

**Figure 4.**
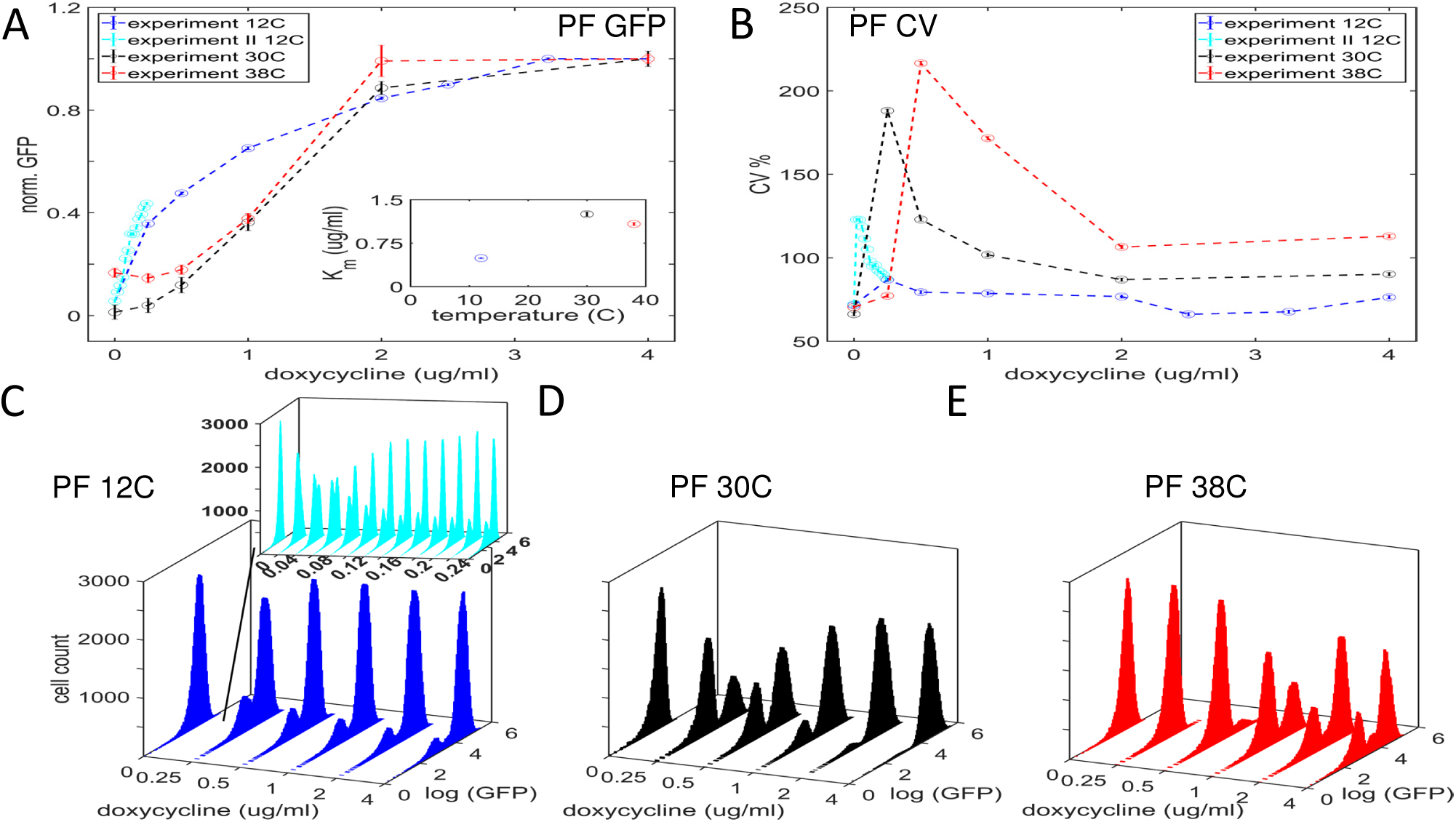
The effect of inducer and temperature on PF strain gene expression dose response. **(A)** Dose-response of the population average (mean) yEGFP::ZeoR expression. Inset shows the doxycycline concentration at which GFP half-saturation (K_m_) occurs as a function of temperature. **(B)** Dose-response of the overall coefficient of variation (CV) of yEGFP::ZeoR expression. Data points from a subsequent dose-response experiment, performed to characterize the CV peak at 12°C are shown in cyan. Error bars are SEM (*N* = 3). **(C)** Fluorescence histograms of yEGFP::ZeoR expression for the PF strain at increasing doxycycline concentrations at 12°C. Inset shows the expression histograms from a subsequent dose response experiment which identified equal peaks at doxycycline = 0.06 μg/ml (axes have the same units as the main figure). **(D)** Fluorescence histograms of yEGFP::ZeoR expression for the PF strain at increasing doxycycline concentrations at 30°C. **(E)** Fluorescence histograms of yEGFP::ZeoR expression for the PF strain at increasing doxycycline concentrations at 38°C.

Cooling and heating preserved the NF gene circuit’s mean dose-response linearity (Nevozhay et al, 2009) nearly up to saturation at all temperatures (Fig. 5A), as predicted computationally (Fig. 3D). Cooling and heating increased the slope of this linear dose response (Fig. 5A), but the effect of cooling was weaker than heating, in contrast to computational predictions (Fig. 3D). Accordingly, both heating and cooling shrank the inducer regime of NF dose-response linearity (Fig. 5A). The inducer-saturated NF expression level dropped in both nonoptimal temperatures (Fig. S6A). Whereas cooling did not alter strongly the flat, uniformly low CV dose-response of NF cells (Fig. 5B), heating had a dramatic effect that contradicted the computational predictions (Fig. 3F and S10D). Remarkably, heating elevated the NF CV dose response to unusually high levels observed only in PF cells (Fig. 4B). Examining the flow cytometry histograms revealed that this surprising CV increase stems from gene expression bimodality of heated NF cells at all levels of induction (Fig. 5E). This contrasted sharply all previous studies of the yeast NF gene circuit (Diao et al, 2016; Nevozhay et al, 2009), where gene expression histograms were always unimodal and narrow, maintaining a low CV at all inducer concentrations (Fig. 5D). Upon closer inspection, we also found a small low-expressing subpopulation for cold NF cells at high induction (Fig. 5C).

**Figure 5.**
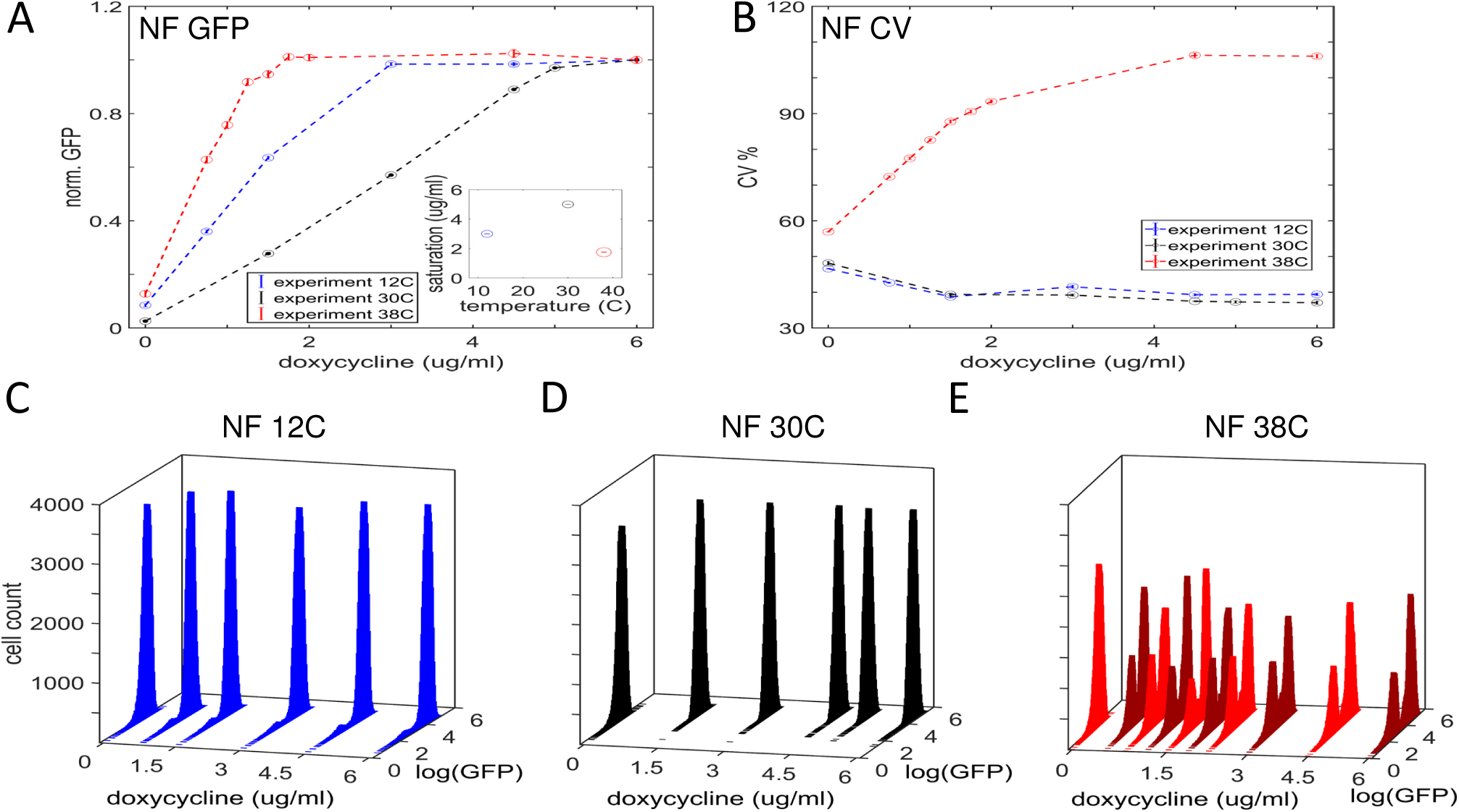
The effect of inducer and temperature on NF strain gene expression dose response. **(A)** Dose-response of the mean of yEGFP::ZeoR expression. Inset shows doxycycline concentration at which GFP expression saturates as a function of temperature. **(B)** Dose-response of the overall coefficient of variation (CV) of yEGFP::ZeoR expression. Error bars are SEM (*N* = 3). **(C)** Fluorescence histograms of yEGFP::ZeoR expression for the NF strain at increasing doxycycline concentrations at 12°C. **(D)** Fluorescence histograms of yEGFP::ZeoR expression for the NF strain at increasing doxycycline concentrations at 30°C. **(E)** Fluorescence histograms of yEGFP::ZeoR expression for the NF strain at increasing doxycycline concentrations at 38°C. For visual clarity, colors alternate between red and maroon.

Overall, the mathematical model predicted reasonably well dose response changes for cold, but not warm NF and PF gene circuits (Figs. 3C-3F and S10). This disagreement between computational predictions and experimental results suggests some reaction-rate independent (non-Arrhenius) effect of heating. Coincidentally, even bacterial (Ratkowsky et al, 1982) and yeast growth (Fig. S2) fails to follow Arrhenius scaling at high temperatures. Therefore, we looked for additional effects of heating that could explain these observations.

### 2.5 Molecular dynamics simulations explain the functional effect of heating

To elucidate the mechanisms underlying the remaining discrepancies between computational and experimental dose responses of heated PF and NF gene circuits, we hypothesized that heating may alter protein structure, affecting transcriptional regulator function. To test this hypothesis, we performed atomistic molecular dynamics (MD) simulations in explicit solvent for the TetR repressor, for which structures are available in the Protein Data Bank (PDB) (see section 4.2 in Materials and Methods, and section 4 in the SI Appendix for details).

TetR functions as a repressor by tightly binding DNA in the major groove, preventing the access of transcriptional machinery. This is possible because the separation of TetR DNA-binding domains (DBDs) is ∼33 Å, which equals the helical pitch of DNA. Previous work (Seidel et al, 2007) has demonstrated that TetR-inducer binding forces the DBDs to move apart, thereby diminishing their propensity to adopt conformations that fit into the major groove of DNA. Therefore, we asked whether temperature might affect the propensity of DNA binding-compatible (or incompatible) TetR DBD conformations.

To address this question, we performed MD simulations on two relevant molecular systems, a TetR:DNA complex (+DNA/-dox; TetR bound to DNA and no doxycycline) and apoTetR (-DNA/+dox; no DNA, but doxycycline is bound to TetR). We simulated these two systems in cold, standard and warm conditions, testing whether the distributions of apoTetR DBD separation distances (Hauser et al, 2016) change with temperature. The results (Fig. 6A) indicate that without inducer the TetR:DNA complex maintains its dynamics at all three temperatures, sampling from a tight distribution (with small standard deviation) centered at a mean value of 33 Å. This matches the helical pitch of DNA, suggesting that temperature does not alter the DNA-binding affinity of uninduced TetR (Hauser et al, 2016). Likewise, cooled apoTetR adopted a large DBD separation distance indicating limited DNA binding propensity compared to standard temperature. However, surprisingly, the DBD separation distance distribution of heated apoTetR shifted to lower values. We verified that this shift is not due to DBD unfolding (Fig. S11) or doxycycline unbinding (Figs. S12 and S13). Unlike for cool and standard temperatures, the warm apoTetR DBD-separation distribution contains the normal TetR:DNA distributions, suggesting that doxycycline-bound apoTetR can abnormally bind to DNA at 38°C. In summary, the MD simulations suggested that heated apoTetR has an increased probability of binding the operator and repressing gene expression despite the presence of inducer.

**Figure 6.**
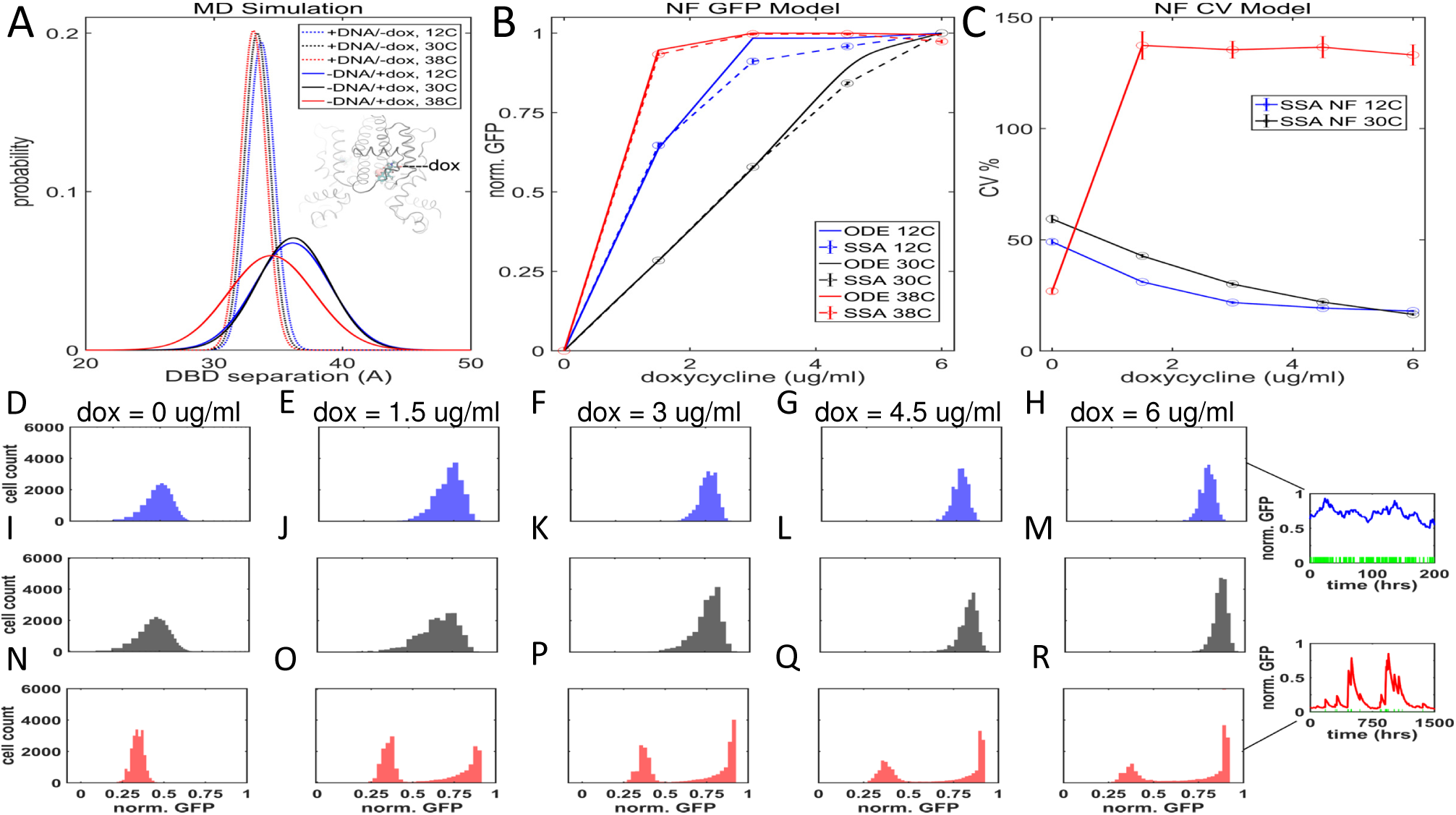
Revised mathematical model based on MD simulation data captures temperature-dependence of dose response and gene expression histograms in NF gene circuit. **(A)** Probability of TetR DNA binding domain (DBD) separation distance as a function of temperature from MD simulations. Distributions of apoTetR with doxycycline bound (-DNA/+dox) and TetR bound to DNA without doxycycline (+DNA/-dox). Inset: Representative structure snapshot at 200 ns (RMSD = 2 Å) from MD simulation with doxycycline docked into the binding pocket of TetR monomer B at T = 38°C. Grey ribbons represent TetR; sticks colored by element (Carbon is cyan)represent Doxycycline; the pink sphere represents the Mg2+ ion. **(B)** Numerical and stochastic dose-response simulations of the mean of yEGFP::ZeoR expression. **(C)** Dose-response of the coefficient of variation (CV) obtained from stochastic simulations of yEGFP::ZeoR expression. Error bars are SEM (*N* = 3). **(D-H)** Stochastically simulated gene expression dose-response histograms corresponding to 12°C. **(I-M)** Stochastically simulated gene expression dose-response histograms corresponding to 30°C, **(N-R)** Stochastically simulated gene expression dose-response histograms corresponding to. All results are represented by blue, black, and red lines and histograms for 12°C, 30°C, and 38°C, respectively. Insets for (H) and (R) show representative promoter kinetics (green bars appear when the P_GAL1-D12_ yEGFP::zeoR promoter is active) and protein dynamics (blue and red lines) for 12°C and 38°C. See Table S2 for parameters.

Motivated by these MD simulation results, we modified the mathematical model to reflect the MD simulation results by allowing DNA binding of apoTetR when heated. To account for DNA-binding compatible and incompatible conformations of apoTetR, we incorporated temperature-dependent terms into the NF Arrhenius model. This revised model captured the increased inducer sensitivity for heated NF gene circuit (Fig. 6B). Stochastic simulations incorporating promoter kinetics (section 3.2 in the SI Appendix) could fully reproduce the experimental heated NF dose response, including the bimodality of expression histograms and correspondingly high CVs (Fig. 6C) (Fig. 6D-6R). Interestingly, gene expression became bimodal only when we lowered the temperature-dependent equilibrium constant *k*_*eq*_, describing promoter kinetics (Fig. 6H,R insets). Therefore, at high temperature the ratio of total TetR (free TetR + apoTetR-dox) DNA-binding (*k*_*b*_) and unbinding (*k*_*u*_) rates must decrease to allow this change in *k*_*eq*_. Indeed, the simulated NF distributions became bimodal only when *k*_*u*_ < *k*_*b*_, suggesting that heating might separately affect both total-TetR unbinding and binding rates to DNA. These results raise new questions about the temperature-dependence of forces underlying protein-DNA binding and their relevance for gene regulatory network fucntion.

Unlike for TetR, there is no structural information in PDB for the rtTA version we employ in the PF gene circuit. Nevertheless, the TetR MD results suggest that the normally high rtTA:doxycycline DNA-binding affinity will diminish at high temperatures. This might explain the lower PF inducer sensitivity and the upward shift of the PF bimodal expression regime.

## 3 Discussion

To mechanistically understand how temperature affects gene regulatory networks, we studied the response of two synthetic gene circuits (PF and NF) to temperature changes. Heating and cooling caused nontrivial, nonmonotone dose-response changes for PF and NF synthetic gene circuits. Despite the noise-suppressing hallmark of negative feedback (Austin et al, 2006; Becskei & Serrano, 2000; Nevozhay et al, 2009), bimodal expression emerged in the NF gene circuit while gene expression noise increased dramatically. Also, the bimodal region shifted upward with temperature for the PF gene circuit.

Growth rates and gene expression can mutually affect each other (Klumpp et al, 2009; Nevozhay et al, 2012; Scott et al, 2010; Tan et al, 2009) and negative feedback has been suggested to offset these effects (Klumpp et al, 2009). By mathematical modeling we demonstrate that, besides growth rate changes, Arrhenius reaction rate scaling is necessary and largely sufficient to explain the effects of cooling, but not those of heating on synthetic gene circuit function. MD simulations suggested that these unexplained effects could result from doxycycline-bound TetR gaining DNA-binding ability in the NF gene circuit, and perhaps a decrease in DNA binding of doxycycline-bound rtTA in the PF gene circuit. Indeed, TetR structure changes gradually as temperature rises up to 40°C (Wagenhofer et al, 1988). Other autoregulators can also gain or lose function depending on temperature (Isaacs et al, 2003). Increased inducer-sensitivities at low temperatures may also result from regulator stabilization, which may alter DNA binding. For example, cooling increases the stability of DNA-bound Arc repressor against pressure denaturation (Foguel & Silva, 1994).

While we investigated synthetic gene circuits, we expect that these conclusions generalize to natural gene regulatory networks as well. Indeed, cooling should generally alter growth rates and reaction rates, which means that cold shock effects on natural gene networks could be predictable before cells attempt to restore homeostasis. In contrast, the effects of heating are probably more complex and harder to predict. Nevertheless, we suggest that MD simulations at high temperatures may provide sufficient information to predict the effects of moderate heating on natural networks. In the future, it will be interesting to investigate how natural negative and positive feedback gene circuits react to heating and cooling, and how these effects alter cellular survival and reproduction in various conditions.

As synthetic biology moves towards real world applications some fundamental challenges will need to be resolved for functional reliability in a diverse range of applications and environments. For example, forward engineering temperature compensation made a synthetic genetic clock robust to temperature changes (Hussain et al, 2014). Inspiration for such designs may come from nature, since temperature compensation appears to be an intrinsic property of natural circadian cycles (Ruoff et al, 1997). Although temperature-induced changes in synthetic gene circuit function present some challenges, they may also present opportunities to exploit this environmental factor for control purposes. For instance, the period of synthetic genetic oscillators can be tuned by altering temperature (Stricker et al, 2008) in addition to inducer levels and growth media. Our findings suggest that achieving a given gene expression level requires less inducer at low temperatures, which may help to reduce the cost of reagents in scientific and industrial applications. At a fundamental level, the insights gained from these models and experiments open new avenues for designing and controlling gene network function and move us towards more fully understanding the dynamics of gene networks in general.

## 4 Materials and Methods

### 4.1 Mathematical modeling

To investigate the effect of temperature dependent growth rate on PF0 and NF0 strain gene expression, we first considered constitutive expression of the yEGFP::ZeoR fluorescent reporter (*z*) according to the model

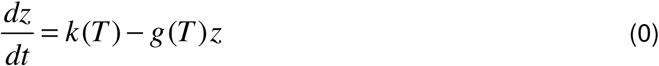

where *k(T) and g(T)* are temperature dependent protein synthesis and growth/dilution rates, respectively.

To quantitatively understand the temperature dependence of the NF and PF gene circuits dose responses, we employed previously developed rate equation models. The equations for the NF gene circuit (Nevozhay et al, 2009) are as follows:

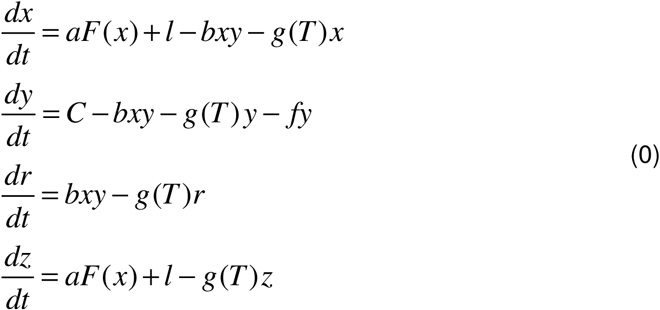

where the variables *x*, *y*, *r*, and *z* denote the intracellular concentration of the inducer-free repressor (TetR) protein, inducer (doxycycline), inducer-bound TetR protein, and the fluorescent reporter (yEGFP::ZeoR) protein concentration. *F(x) = θ*^*n*^ /(*θ*^*n*^ *+ x*^*n*^), where *θ* is the repression threshold corresponding to an effective repressor-DNA dissociation constant and *n* is the Hill coefficient. *C* is a control parameter that describes the rate of doxycycline entry into the cell and is proportional to extracellular inducer concentration. The repressor and reporter-resistance proteins are synthesized at a rate *aF*(*x*) (same rate constant *a* as both genes are regulated by the same P_GAL1-D12_ promoter). Dilution due to temperature dependent cellular growth of all three variables is *g*(*T*); the inducer also degrades at a rate *f* (the degradation of TetR and yEGFP::ZeoR have been neglected due to their relatively long timescales compared to the degradation of the inducer and cell division time in yeast (Nevozhay et al, 2009). The leaky protein synthesis rate is *l* and the inducer-repressor binding rate is *b*.

Based on the results of the MD simulations (Section 2.5), we modified the NF gene circuit model (Eqn. 2) to incorporate a temperature-dependent parameter *m*(*T*) that describes the fraction of apoTetR with a conformational state conducive P_GAL1-D12_ promoter binding. This modified model is identical to the model presented in Equation 2 except *F*(*x*) becomes *F*(*x+mr*) *= θ*^*n*^ /[*θ*^*n*^ *+* (*x + mr*)^*n*^]. Details for a stochastic implementation of this modified model are in Section S3.1, and a full stochastic model that incorporates promoter dynamics in Section S3.2.

The PF gene circuit can be described by the following equations (Nevozhay et al, 2012):

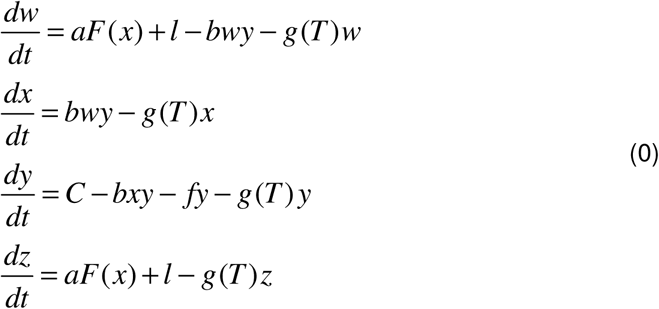

where *w*, *x*, *y*, and *z* correspond to inactive rtTA, active rtTA, inducer, reporter/resistance protein concentrations, respectively. Here, *F(x) = x*^*n*^*/(θ*^*n*^ *+ x*^*n*^) where *θ* is the induction threshold corresponding to an effective activator-DNA dissociation constant. All other parameters are as described for Equation 2.

We obtained the parameters for the 30°C condition by fitting the PF and NF models to the experimental data using custom Matlab scripts to minimize the sum of squared errors, starting from parameters obtained from (Nevozhay et al, 2009; Nevozhay et al, 2012). We set the dilution rate *g* for each temperature equal to the mean cellular growth rate obtained from exponential fits to the experimental growth rate data shown in Figure S7A,B. Note that the Hill function constants have been considered temperature independent, as they are ratios of rate constants and are specific for a certain reaction (Heiland et al, 2012). The Arrhenius scaling of reaction rates for the 12°C and 38°C temperature conditions are discussed in section 2 in SI Appendix.

### 4.2 Molecular dynamics simulations

We based the model of TetR (class D) bound to DNA on the X-ray crystal structure PDB ID: 1QPI (Orth et al, 2000). We used the biological unit to obtain the homodimer:DNA complex. Each monomer of TetR was missing an eight-residue loop (between Leu155 and Glu164). We reconstituted this missing loop using a homology model subsequently relaxed using minimization and MD, merging the loop with TetR without adding strain (Fig. S11). The inducer binding pocket was occupied by imidazole. We thus removed imidazole and docked doxycycline: Mg^2^+ instead by merging the coordinates with a second TetR structure [PDB ID: 1TRT (Hinrichs et al, 1994)] containing Doxycycline: Mg^2^+ (see section 4.3 in SI Appendix). We added three base pairs of GCG to the termini of the DNA to reduce end-fraying (Galindo-Murillo et al, 2014) by RMS-aligning the phosphate linkages to those of the DNA in the above model and merging the coordinates. This resulted in a holo TetR model (+DNA/+dox, i.e. bound to DNA, “+DNA” and bound to doxycycline, “+dox”). The apoTetR model (-DNA/+dox) was the result of unmerging DNA from the above model; the holo TetR model (+DNA/-dox) was the result of unmerging dox from the above model. A truncated octahedron with 10 Å buffer of explicit water and (0.2 M) KCl ions solvated these two systems. We used TIP3P (Jorgensen et al, 1983) parameters for water, Joung-Cheatham (Joung & Cheatham, 2008) parameters for K^+^ Cl^-^ ions, Li et al. (Li et al, 2013) parameters for Mg^2^+, ff99SB parameters (Hornak et al, 2006) for protein and parmBSC0 parameters (Perez et al, 2007) for DNA (Hauser et al, 2016). We used an 8 Å non-bonded cutoff; and PME (Essmann et al, 1995) to calculate long-range electrostatics. We performed equilibration using a previous ten stage approach (Hauser et al, 2016) (Table S1), with three independent runs at three temperatures (12°C, 30°C and 38°C). After equilibration, we performed 200 ns of production MD (3.6 μs total MD). Based on previous work with MTERF1 (Hauser et al, 2016), we obtained TetR superhelical pitch from the coordinates of the Cα atoms of conserved Pro residues tracking the DNA major groove. TetR superhelical pitch is equivalent to the linear distance between Pro36 (DBD of monomer A) and Pro239 (DBD of monomer B).

### 4.3 Strains and media

The haploid Saccharomyces cerevisiae strain YPH500 (*α*, *ura3-52*, *lys2-801*, *ade2-101*, *trp1Δ63*, *his3Δ200*, *leu2Δ1*) (Stratagene, La Jolla, CA) was used as a model organism throughout this study. Cultures were grown in synthetic drop-out medium with the appropriate supplements (SD -his -trp +ade) to maintain selection and supplemented with sugars (glucose or galactose) as described below (all reagents from Sigma, St. Louis, MO). Doxycycline (Fisher Scientific, Fair Lawn, NJ) was used to induce the PF and NF gene circuits.

### 4.4 Cell culture

Well-isolated single yeast colonies were picked from plates and incubated overnight in synthetic drop-out medium supplemented with 2% glucose at 30°C. Twelve hours later, the 1 ml cell suspensions were diluted to the concentration 1 × 10^6^ cells/ml -concentration estimated using a Nexcelom Cellometer Vision cell counter (Nexcelom Bioscience, Lawrence, MA) - in fresh medium supplemented with 2% galactose. Triplicate cultures were grown for 48 hours in galactose medium with the appropriate concentration of doxycycline (Acros Organics, Geel, Belgium) for each condition to allow gene expression levels to stabilize (Nevozhay et al, 2009; Nevozhay et al, 2012). Cell density was measured (Nexcelom Cellometer Vision) and resuspended to a concentration 1×10^6^ cells/ml every 12 or 24 hours depending on the experimental condition to keep them in log growth phase.

### 4.5 Flow cytometry

Population gene expression was read on the BD Accuri flow cytometer (Becton Dickinson, Franklin Lakes, NJ) after 48 hours of doxycycline induction. We considered gene expression being stable when GFP distributions for 30°C and 38°C experiments did not change between 24 hour and 48 hour measurements (PF cells: Figs. 4D,E and S14B,C; NF cells: Figs. 5D,E and S14E,F), and similarly, the distributions for 12°C did not change between 48 hour and 72 hour measurements (PF cells: Figs. 4C and S15A; NF cells: Figs. 5C and S15B).

### 4.6 Fitness measurements

Malthusian fitness between subsequent resuspensions was estimated by linear fits to log-transformed cell counts (inferred from cell density measurements described in Section 4.4 and culture volume).

### 4.7 Data processing and analysis

Raw flow cytometry data files were read into Matlab (Mathworks, Inc.) using the Matlab script fca readfcs (Matlab Central) for plotting and analysis. A small gate was applied to the forward scatter and side scatter data to minimize the contribution of extrinsic noise due to cell cycle phase, cell size and age (Newman et al, 2006), and exclude doublets, dead cells, and cellular debris from our analysis. To eliminate small numbers of mutated cells that may have lost the integrated construct (due to homologous recombination) or rare cells left over from previous samples (not eliminated by flow cytometer), cells with log fluorescence deviating more than 3 standard deviations from the mean were considered outliers and discarded from the analysis (Diao et al, 2016; Nevozhay et al, 2009; Nevozhay et al, 2012).

## Acknowledgments

This research was supported by an NSERC Postdoctoral Fellowship (PDF-453977-2014) to D.C., NIH NRSA Fellowship (F31-GM101946) and NSF AGEP-T Fellowship (HRD-1311318) to K.H., NIH MIRA grant (R35GM122561) to G.B., and the Laufer Center for Physical and Quantitative Biology.

## Author contributions

D.C. and G.B. designed the research, D.C. and S.M. performed the experiments, D.C., G.B. and K.H. performed the mathematical modeling and MD simulations, D.C. analyzed the data, and D.C., K.H., and G.B. wrote the paper.

